# Transferable IncX3-*bla*_NDM-15_ in an uncommon ST580 *Klebsiella pneumoniae* recovered during paediatric intensive-care surveillance

**DOI:** 10.64898/2026.08.11.744171

**Authors:** Zhenghao Lou, Chengcheng Ye, Xiaolu Yang, Qiyu Liu, Chennuo Wang, Hao Xu, Beiwen Zheng, Xiawei Jiang

**Author notes:** Corresponding authors: Xiawei Jiang, Email address, Address: Zhejiang Chinese Medical University, No. 548 Binwen Road, Binjiang District, Hangzhou City, Zhejiang Province, China (310053). These authors contributed equally to this work and share first authorship (Zhenghao Lou, Chengcheng Ye). These authors contributed equally to this work and share corresponding authorship (Beiwen Zheng, Xiawei Jiang).

## Abstract

**Objective:** Carbapenem-resistant *Klebsiella pneumoniae* harboring *bla*_NDM_ poses a serious threat to public health; however, *bla*_NDM-15_ remains poorly characterized outside the dominant epidemic lineages.

**Methods:** We characterized *K. pneumoniae* strain ETFK6090, isolated from a perianal surveillance swab of an 11-month-old immunocompromised child in a paediatric intensive care unit. Investigations included antimicrobial susceptibility testing, broth conjugation, S1 nuclease PFGE with Southern blotting, complete genome sequencing, and comparative genomic analysis against 465 curated *bla*_NDM_-positive *K. pneumoniae* genomes from 37 countries.

**Results:** ETFK6090 belonged to ST580 and exhibited resistance to carbapenems, ceftazidime-avibactam, broad-spectrum cephalosporins, fluoroquinolones, gentamicin, chloramphenicol and trimethoprim-sulfamethoxazole; amikacin and fosfomycin retained low MICs. The complete genome comprised one chromosome and five plasmids, *bla*_NDM-15_ was localized on a 46,161-bp IncX3 plasmid, confirmed by Southern blotting. Conjugation into *Escherichia coli* EC600 transferred carbapenem and cephalosporin resistance, confirming in vitro mobility. The *bla*_NDM-15_ genetic environment retained a conserved *bla*_NDM_ module, with IS-mediated rearrangements at the downstream boundary. In the global comparison, *bla*_NDM-1_ and *bla*_NDM-5_ predominated, the ST580-*bla*_NDM-15_ combination was exceedingly rare, and ETFK6090 constituted a distinct branch apart from major epidemic lineages.

**Conclusions:** A transferable IncX3-*bla*_NDM-15_ plasmid can emerge in an uncommon ST580 background, underscoring the necessity to extend genomic surveillance of carbapenem-resistant *K. pneumoniae* beyond dominant epidemic clones, particularly in high-risk paediatric and intensive-care settings.

## 1. Introduction

Carbapenem-resistant *Klebsiella pneumoniae* (CRKP) remains a major challenge for clinical management and infection control because treatment options are limited, attributable mortality is substantial and mobile resistance determinants disseminate efficiently Among the mechanisms underlying carbapenem resistance (1, 2), carbapenemase production is particularly important, and the global spread of New Delhi metallo-beta-lactamase (NDM) has received sustained attention (3). *bla*_NDM-1_ was first described in 2008 in a *K. pneumoniae* isolate from a Swedish patient of Indian origin in New Delhi (4). Since then, the *bla*_NDM_ family has continued to diversify into multiple variants with distinct epidemiological and functional features.

*bla*_NDM-15_ is an infrequently reported NDM allele that differs from *bla*_NDM-1_ by two amino acid substitutions, M154L and A233V. Experimental work on NDM variants suggests that substitutions at these positions may influence enzyme stability under zinc-restricted conditions and may therefore provide an adaptive advantage in specific host environments (5, 6). However, the epidemiological settings in which *bla*_NDM-15_ circulates remain incompletely defined. In particular, information on its plasmid backbone, local genetic environment, host clonal background and position within the broader *bla*_NDM_-positive *K. pneumoniae* population is limited.

Here, we describe ETFK6090, a *bla*_NDM-15_-positive *K. pneumoniae* isolate recovered from an 11-month-old child in a paediatric intensive care unit (PICU). The patient was receiving intensive cytotoxic chemotherapy under the HLH-1994 protocol for haemophagocytic lymphohistiocytosis and was severely immunosuppressed with neutropenia. We combined phenotypic testing, conjugation assays, plasmid localization, complete genome sequencing and comparative genomics to characterize the isolate, define the genetic context and transferability of its IncX3 plasmid, and place it within the phylogenetic context of publicly available complete *bla*_NDM-_positive *K. pneumoniae* genomes. Because the isolate was recovered during colonization screening rather than from invasive infection, its clinical relevance is interpreted primarily in terms of colonization, silent carriage and transmission risk.

## 2. Materials and Methods

### 2.1 Strain isolation and identification

*K. pneumoniae* ETFK6090 was isolated in June 2022 from a perianal swab specimen obtained from an 11-month-old child in the PICU of the First Affiliated Hospital of Zhengzhou University, China. The sample was inoculated into brain heart infusion (BHI) broth and incubated with shaking at 37°C for 18–24 h. The culture was then diluted with phosphate-buffered saline and spread onto Columbia blood agar plates, followed by incubation at 37°C for 24 h. Distinct colonies were selected according to colony morphology and subcultured to purity. Species identification was performed using matrix-assisted laser desorption/ionization time-of-flight mass spectrometry (MALDI-TOF MS).

### 2.2 Antimicrobial susceptibility testing

Minimum inhibitory concentrations (MICs) of 21 antimicrobial agents were determined using agar dilution or broth microdilution, as appropriate. Susceptibility to tigecycline and polymyxin B was interpreted according to European Committee on Antimicrobial Susceptibility Testing (EUCAST) clinical breakpoints (v15.0). Omadacycline and eravacycline were interpreted according to United States Food and Drug Administration criteria. All other agents were interpreted using Clinical and Laboratory Standards Institute (CLSI) M100-Ed35 criteria. *Escherichia coli* ATCC 25922 and *K. pneumoniae* ATCC 700603 were used as quality-control strains.

### 2.3 Whole-genome sequencing and analysis

Genomic DNA was extracted using a commercial DNA extraction kit (QIAGEN, Hilden, Germany). Whole-genome sequencing was performed on an Oxford Nanopore platform, and raw reads were assembled into a complete genome using Unicycler (7). Assembly quality was assessed using CheckM (8), and genome annotation was performed with Prokka (9). Acquired antimicrobial resistance genes were identified using the Comprehensive Antibiotic Resistance Database (CARD). Virulence factors were identified by BLAST searches against the Virulence Factor Database. Insertion sequences and transposable elements were detected using ISFinder. Multilocus sequence typing (MLST) was conducted using the Institut Pasteur *Klebsiella* database (http://bigsdb.web.pasteur.fr/klebsiella/klebsiella.html), and plasmid replicon types were determined using PlasmidFinder together with mlplasmids.

### 2.4 Conjugation assay

The transferability of the *bla*_NDM-15_ plasmid was evaluated by broth conjugation, using ETFK6090 as the donor and rifampicin-resistant *E. coli* EC600 as the recipient. Donor and recipient cultures in logarithmic growth phase were mixed at a 2:1 volume ratio in LB broth and incubated at 37°C for 18–24 h. Serial dilutions were plated onto LB agar containing rifampicin (600 mg/L) and imipenem (2 mg/L). Putative transconjugants were purified and verified by MALDI-TOF MS and PCR amplification of *bla*_NDM_.

### 2.5 S1-PFGE and Southern blotting

Plasmid size and *bla*_NDM-15_ localization were analysed by S1 nuclease pulsed-field gel electrophoresis (S1-PFGE) followed by Southern blotting. ETFK6090 bacterial suspensions were embedded in agarose plugs and digested with S1 nuclease at 37°C for 30 min. PFGE was performed using a CHEF-Mapper XA system (Bio-Rad, USA) at 300 k for 16 h. *Xba*I-digested *Salmonella* enterica serovar Braenderup H9812 was used as the molecular-weight marker. DNA was transferred to a positively charged nylon membrane and hybridized with a DIG-labelled *bla*_NDM-15_-specific probe. Signals were detected using the DIG DNA Labelling and Detection Kit (Roche).

### 2.6 Comparative genomic analysis

The genetic environment surrounding *bla*_NDM-15_ and the backbones of related plasmids were visualized using Easyfig, and comparative plasmid analysis was performed with Proksee. A total of 5,246 complete *bla*_NDM_-carrying genome sequences were retrieved from the NCBI Pathogen Detection platform on 24 November 2025. After restriction to *K. pneumoniae* and application of dereplication and quality-control procedures, 465 complete *bla*_NDM_-positive genomes from 37 countries, including ETFK6090, were retained for comparative analysis. Only *bla*_NDM_ annotations labelled as complete were included in subtype counts. Core-gene analysis was performed with Roary (10), and a maximum-likelihood phylogeny was inferred using IQ-TREE and visualized in iTOL (11). To compare genomes carrying the two predominant subtypes, *bla*_NDM-1_ and *bla*_NDM-5_, we first performed univariable analyses and then fitted a binary logistic regression model to identify factors independently associated with subtype status. Categorical variables were compared using Pearson’s chi-square test or Fisher’s exact test, as appropriate. Skewed continuous variables were analysed using the Mann– Whitney U test. Variables with *p* < 0.10 in univariable analyses were entered into the multivariable model (https://proksee.ca/ https://itol.embl.de/).

### 2.7 Structural prediction and molecular docking analysis

The amino acid sequence encoded by *bla*_NDM-15_ was inferred from the assembled gene and aligned with reference NDM variants. The three-dimensional structure of NDM-15 was predicted using AlphaFold, evaluated using model-confidence metrics and visualized in PyMOL to compare key residues and local conformational features (12). Molecular docking with relevant antimicrobial ligands was performed using AutoDock Vina after standard receptor and ligand preparation. Binding energies, top-ranked poses and predicted protein–ligand interactions were analysed to explore potential structural and substrate-binding effects of the NDM-15-associated substitutions. These analyses were considered exploratory and were not used to infer quantitative enzymatic activity.

### 2.8 STORMS Checklist

This study has been reported in accordance with the Strengthening The Organization and Reporting of Microbiome Studies (STORMS) checklist (version 1.03) (13). The completed checklist has been deposited in Zenodo and is accessible at https://doi.org/10.5281/zenodo.21866628.

## 3 Results

### 3.1 Clinical context, antimicrobial susceptibility profile and transferability

K. *pneumoniae* ETFK6090 was recovered from a perianal surveillance swab from an immunocompromised 11-month-old child in the PICU and showed an extensively drug-resistant phenotype. High MICs were observed for carbapenems (imipenem and meropenem), broad-spectrum cephalosporins, ceftazidime-avibactam, fluoroquinolones, gentamicin, chloramphenicol, trimethoprim-sulfamethoxazole, omadacycline and eravacycline, whereas amikacin and fosfomycin retained low MICs (Table S1). Broth conjugation yielded a transconjugant, designated 6090EC. Compared with the recipient background, 6090EC acquired markedly elevated MICs to amoxicillin-clavulanate, piperacillin-tazobactam, third- and fourth-generation cephalosporins, ceftazidime-avibactam, imipenem and meropenem, whereas most non-beta-lactam MICs decreased substantially. These data support the in vitro transferability of the principal beta-lactam resistance determinant and indicate that the broader multidrug-resistant phenotype of ETFK6090 was only partly explained by the conjugative element.

### 3.2 Complete genome features and localization of *bla*_NDM-15_ on an IncX3 plasmid

Complete genome sequencing assigned ETFK6090 to ST580 and resolved one circular chromosome and five circular plasmids (Table S2, Figure 1). Three plasmids were typeable as IncN (247,777 bp), IncY (99,157 bp) and IncX3 (46,161 bp), whereas two smaller plasmids were untypeable. Resistome analysis identified 22 acquired resistance genes in addition to the chromosome-associated genes *fosA* and *oqxAB*. *bla*_NDM-15_ was located on the IncX3 plasmid, whereas *bla*_DHA-1_, *bla*_TEM-1B_ and *qnrB4* were located on the IncN plasmid. S1-PFGE resolved three major plasmid bands of approximately 750– 850 kb, 200–215 kb and 40–50 kb (Figure 2a). The difference between the PFGE profile and the complete assembly most likely reflected the limited resolution of small plasmids by S1-PFGE. Southern blotting with a *bla*_NDM-15_ probe produced a single signal on the 40–50-kb band (Figure 2a), consistent with localization of *bla*_NDM-15_ on the 46,161-bp IncX3 plasmid.

**Figure 1.**
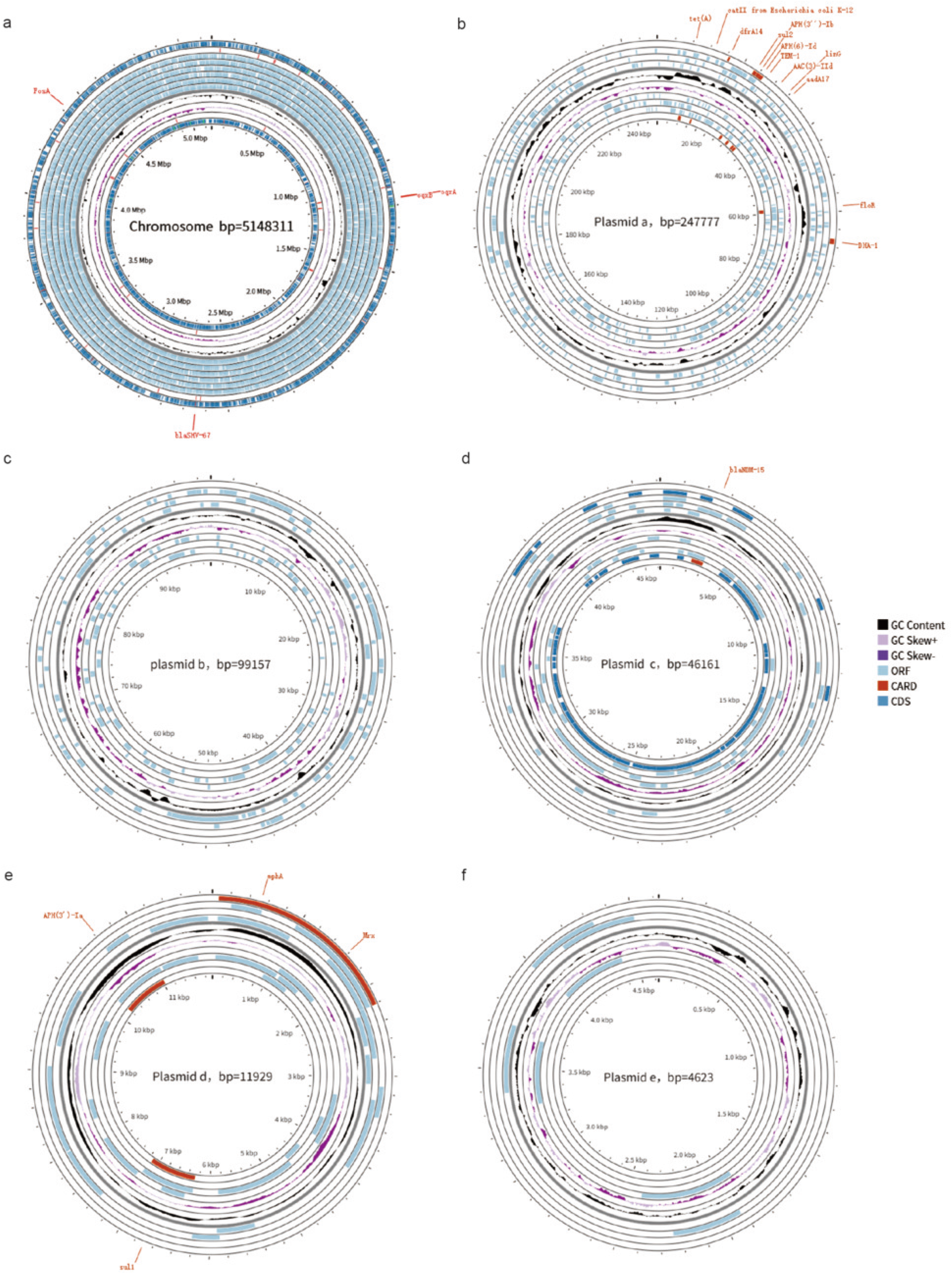
Genomic map of *K. pneumoniae* ETFK6090. **(a)** shows the circular chromosome, and panels **(b–f)** show the five plasmids resolved by long-read assembly. Each plot displays, from outside to inside, forward-strand CDS, reverse-strand CDS, antimicrobial resistance genes annotated by CARD, ORFs, GC skew and GC content.

**Figure 2.**
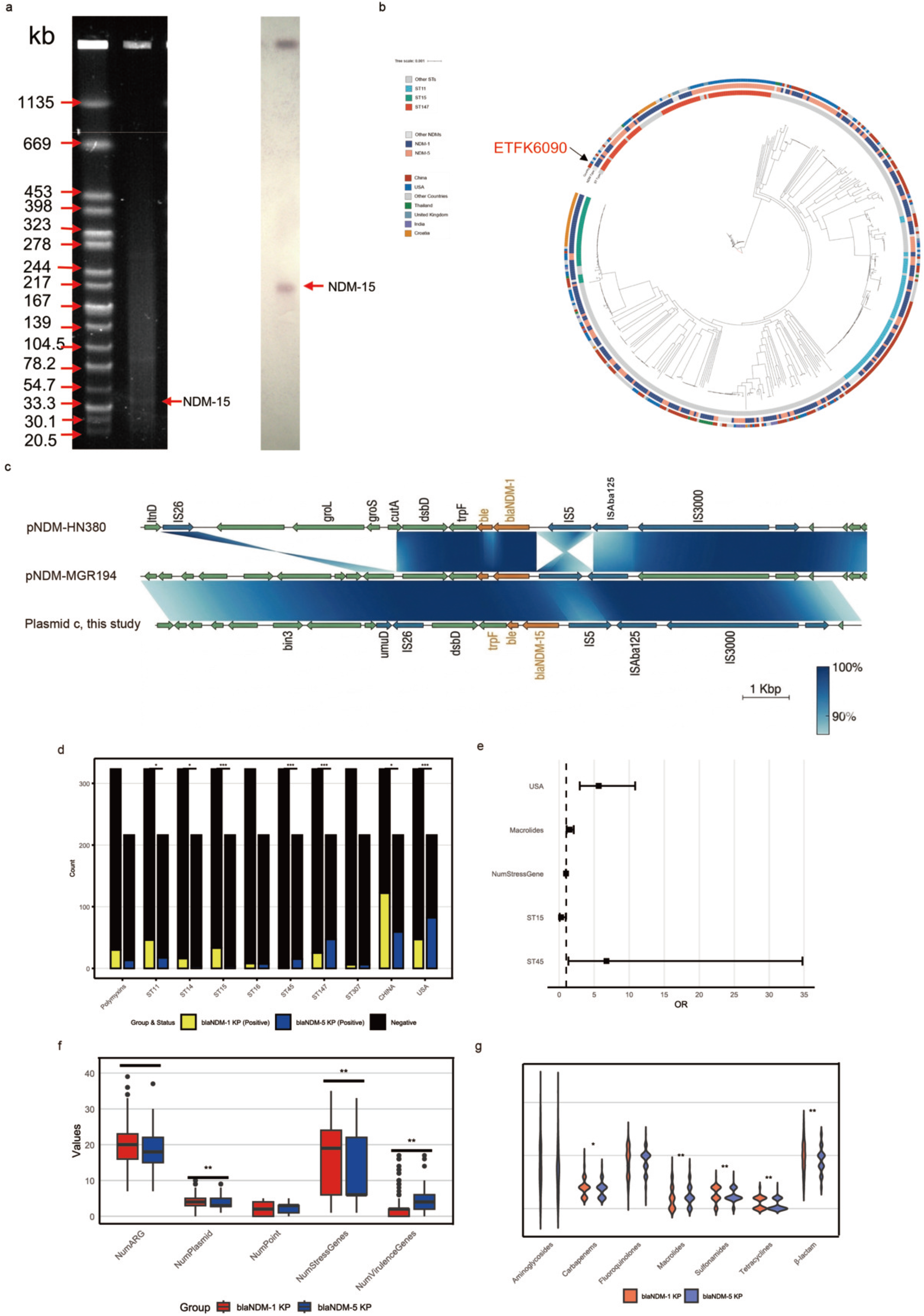
Genomic characterization of ETFK6090 and phenotypic/genotypic comparison of *bla*_NDM_-positive *K. pneumoniae*. **(a)** Localization of the *bla*_NDM-15_ plasmid in ETFK6090 by S1 nuclease pulsed-field gel electrophoresis (left) and Southern blotting with a *bla*_NDM-15_-specific probe (right). **(b)** Core-gene phylogeny of 465 *bla*_NDM_-positive *K. pneumoniae*. The maximum-likelihood tree shows the relationship between ETFK6090 and the global comparison set. **(c)** Linear comparison of the genetic environment of *bla*_NDM-15_ in ETFK6090. The upper sequences represent the reference *bla*_NDM_-associated regions, and the lower sequence represents the corresponding *bl*a_NDM-15_ region in ETFK6090. Arrows indicate ORFs and transcription direction; orange indicates antimicrobial resistance genes, dark blue indicates mobile genetic elements, and green indicates other genes. Blue shading indicates sequence homology. Compared with the reference sequence, the cutA-groS-groL module is absent in ETFK6090 and is replaced by sequences adjacent to IS26. d–g: Comparison of *bla*_NDM-1_- and *bla*_NDM-5_-positive *K. pneumoniae*. **(d)** Distribution across major sequence types and geographical origins. **(e)** Multivariable logistic regression results, *K. pneumoniae* harbouring *bla*_NDM-1_ served as the reference group. **(f)** Comparison of resistance-gene counts, plasmid replicons, point mutations, stress genes, and virulence genes. **(g)** Distribution of acquired resistance genes by antibiotic class.

### 3.3 Genetic environment of *bla*_NDM-15_ in ETFK6090

Comparison of sequences flanking *bla*_NDM-15_ showed that the core resistance region in ETFK6090 was highly similar to reference *bla*_NDM_-associated contexts (Figure 2c). The downstream *ble-trpF-dsbD* cluster was intact, and the upstream region retained an IS*5*-truncated IS*Aba125*-associated structure typical of many *bla*_NDM_ environments. The main structural difference was located downstream of *dsbD*. Whereas the reference sequence was linked to the *cutA-groS-groL* module, ETFK6090 carried IS*26*, *umuD* and *bin3* in this region, suggesting insertion-sequence-mediated rearrangement at the downstream boundary. These findings indicate that the *bla*_NDM-15_ module in ETFK6090 is broadly conserved but exhibits localized structural variability at its margins.

### 3.4 Comparative genomic context and phylogenetic position of ETFK6090 among *bla*_NDM_-positive *K. pneumoniae*

Among the 465 curated complete *bla*_NDM_-positive *K. pneumoniae* genomes from 37 countries collected between 2009 and 2025, *bla*_NDM-1_ and *bla*_NDM-5_ were the dominant subtypes, accounting for approximately 60% and 34% of genomes, respectively. *bla*_NDM-1_ showed a broader geographical distribution, particularly in China, the United States and Croatia, whereas *bla*_NDM-5_ was concentrated mainly in China and the United States (Figure 3). The geographical distributions differed significantly (*χ*² = 38.97, *p* < 0.001), and genomes from the United States were more likely to carry *bla*_NDM-5_ (aOR = 5.64, 95% CI: 2.93–10.85, *p* < 0.001). The two subtype groups also differed in clonal background. *bla*_NDM-1_ was mainly associated with ST11 and ST15, whereas *bla*_NDM-5_ was enriched in ST147 (Figure 2d-g). Multivariable analysis indicated that ST15 (OR = 0.26, 95% CI: 0.07–0.93, *p* = 0.039) were associated with *bla*_NDM-1_, while ST11 (*p* = 0.828) and ST147 (*p* = 0.073) showed no significant association with a specific subtype.

**Figure 3.**
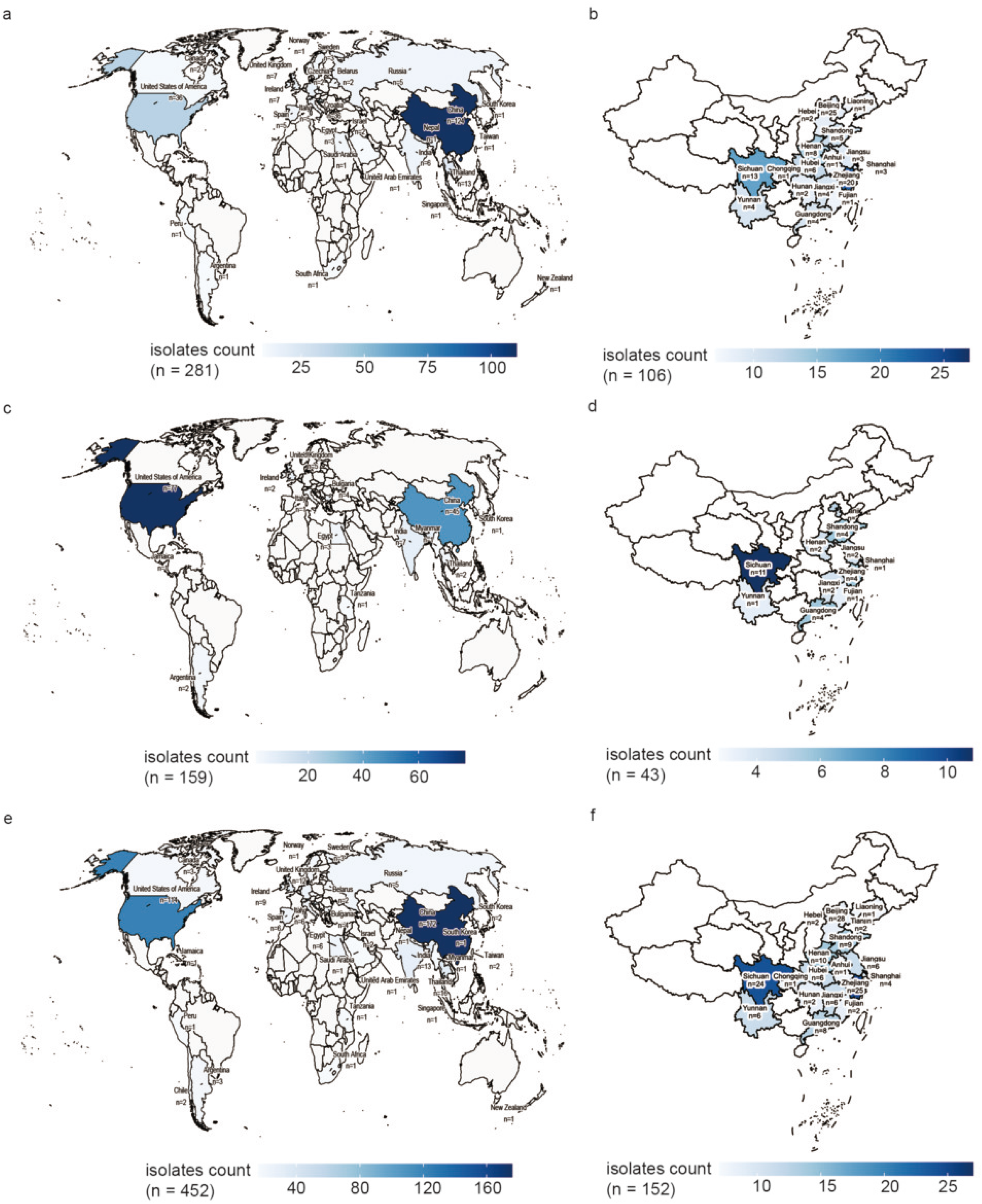
Geographical distribution of *bla*_NDM_-carrying *K. pneumoniae* genomes included in the curated comparative dataset. **(a)** Global distribution of *bla*_NDM-1_; **(b)** Chinese distribution of *bla*_NDM-1_; **(c)** Global distribution of *bla*_NDM-5_; **(d)** Chinese distribution of *bla*_NDM-5_; **(e)** Global distribution of all *bla*_NDM_ variants; **(f)** Chinese distribution of all *bla*_NDM_ variants. Colour intensity indicates the number of genomes recovered from each location.

In univariable comparisons, *bla*_NDM-5_-positive genomes showed greater plasmid-replicon complexity and higher numbers of acquired resistance, virulence and stress-related genes (Figure 2d-g). After adjustment, however, only increased macrolide resistance-gene abundance and reduced stress-gene abundance remained significant. These results suggest that some apparent subtype-associated genomic differences are partly explained by clonal background and geographical structure.

Within this global dataset, ETFK6090 belonged to ST580, an uncommon lineage relative to the predominant *bla*_NDM_-associated sequence types. In the core-gene phylogeny, ETFK6090 formed a distinct branch separated from the major ST11 and ST147 clusters (Figure 2b), indicating that *bla*_NDM-15_ in this isolate occurred in a genetic background distinct from the dominant global *bla*_NDM_-positive *K. pneumoniae* lineages.

### 3.5 Exploratory structural modelling and molecular docking analysis of NDM-15

ETFK6090 carried a full-length 813-bp *bla*_NDM-15_ gene that shared 99.75% identity with *bla*_NDM-1_ and encoded two amino acid substitutions, M154L and A233V. Structural superposition with the NDM-1 crystal structure (PDB: 4EYF) showed a highly conserved overall backbone, with both substitutions located outside the catalytic core (Figure 4a–c) (14). Docking analysis indicated that meropenem could adopt a stable predicted binding pose in the NDM-15 active pocket, with a binding energy of −6.058 kcal/mol, involving the catalytic zinc ion and the conserved residues Asn220 and Asp124 (Figure 4d, e). These exploratory results support a conserved active-pocket architecture in NDM-15, while suggesting possible local structural effects of the distal substitutions.

**Figure 4.**
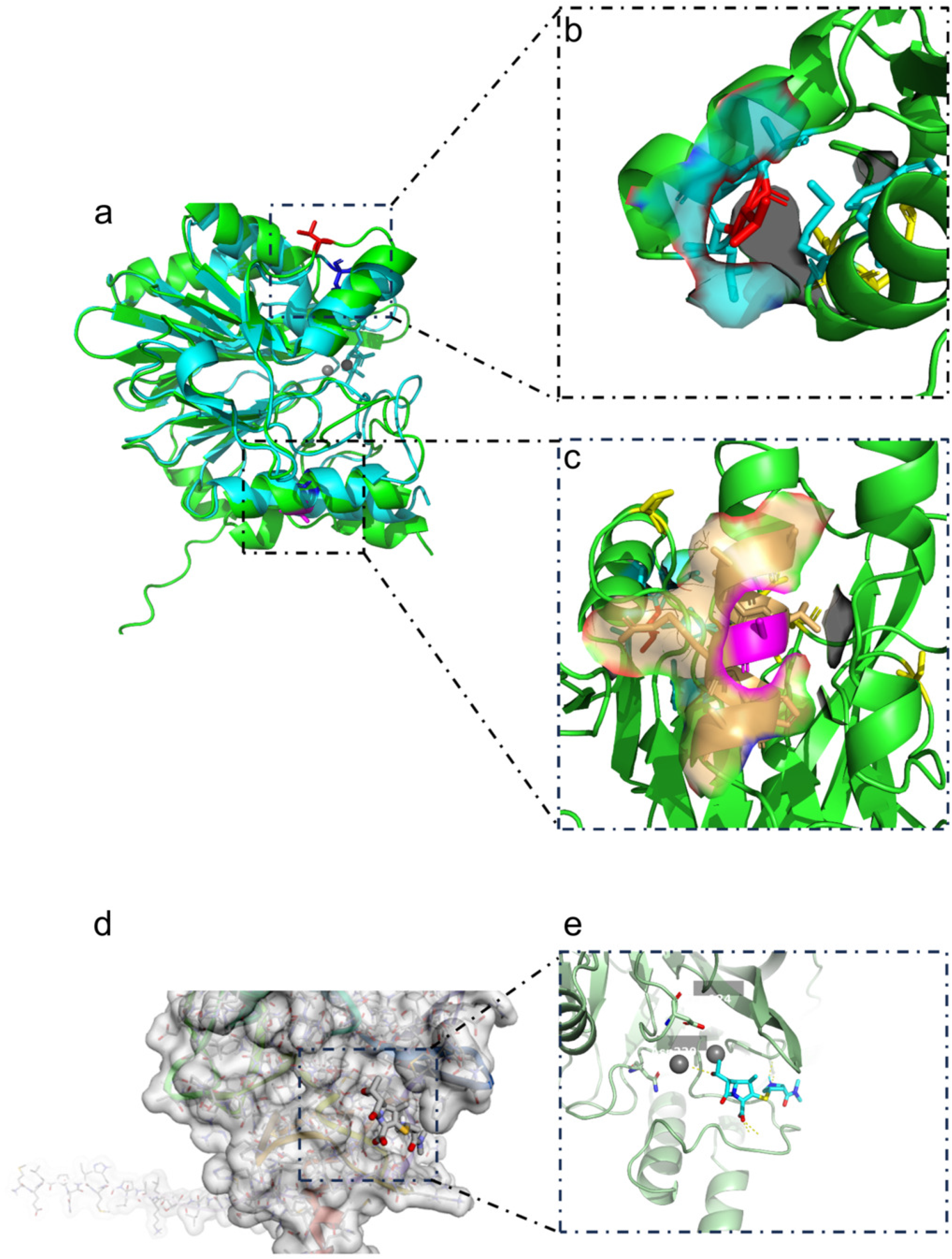
Exploratory structural modelling of NDM-15. **(a)** Superposition of the AlphaFold-generated NDM-15 model and the reference NDM-1 structure. **(b, c)** Local views of the M154L and A233V substitutions. **(d)** Docking pose of meropenem in the predicted active-site pocket. **(e)** Schematic view of local interactions involving conserved active-site residues. These analyses are exploratory and were not used to infer quantitative enzymatic activity.

## 4 Discussion

In this study, we characterized an extensively drug-resistant *K. pneumoniae* isolate, ETFK6090, recovered during paediatric intensive-care colonization screening. The principal carbapenem resistance determinant, *bla*_NDM-15_, was carried by a 46,161-bp IncX3 plasmid that was transferable in vitro. Importantly, the isolate belonged to ST580 rather than to the major *bla*_NDM_-associated epidemic lineages represented in our curated complete-genome dataset, and the resistance locus was embedded within a conserved *bla*_NDM_-associated mobile module. Together, these findings indicate that clinically important *bla*_NDM_ dissemination is not restricted to dominant epidemic clones and can occur in less common genomic backgrounds (15).

The resistance phenotype of ETFK6090 is notable because carbapenem resistance was accompanied by resistance to ceftazidime-avibactam and multiple non-beta-lactam classes, leaving few agents with retained in vitro activity. The conjugation experiments clarified the genetic basis of this phenotype. Transfer to *E. coli* EC600 reproduced the major beta-lactam resistance profile, including resistance to carbapenems and broad-spectrum cephalosporins, whereas most non-beta-lactam resistance phenotypes were not co-transferred. This pattern is consistent with a transferable plasmid carrying the principal beta-lactam resistance determinant within a broader multicomponent resistance background. Complete-genome analysis supports this interpretation: *bla*_NDM-15_ was assigned to the IncX3 plasmid, whereas additional acquired determinants, including *bla*_DHA-1_, *bla*_TEM-1B_ and *qnrB4*, were located on a larger IncN plasmid. IncX3 backbones have repeatedly been implicated in *bla*_NDM_ dissemination among Enterobacterales, and our findings reinforce the continuing epidemiological relevance of this plasmid type in mediating transferable carbapenem resistance (16).

The *bla*_NDM-15_ gene in ETFK6090 encodes the M154L and A233V substitutions relative to NDM-1. Our modelling suggests that these substitutions do not disrupt the overall active-pocket architecture, which is consistent with the preserved carbapenem-resistant phenotype. However, because AlphaFold modelling and molecular docking cannot substitute for enzyme kinetics or zinc-depletion experiments, these structural data should be interpreted as hypothesis-generating rather than definitive functional evidence. The revised interpretation is therefore conservative: the NDM-15 substitutions may influence local conformation or stability while preserving the core hydrolytic architecture, but their functional consequences require biochemical validation (17).

The comparative genomic analysis places ETFK6090 in a broader surveillance context. Within the curated set of 465 complete *bla*_NDM_-positive *K. pneumoniae* genomes, *bla*_NDM-1_ and *bla*_NDM-5_ predominated and were associated mainly with established sequence types such as ST11, ST15 and ST147. In contrast, the ST580-*bla*_NDM-15_ combination was uncommon, and ETFK6090 formed a distinct phylogenetic branch outside the major epidemic clusters. These data suggest that *bla*_NDM-15_ in ETFK6090 was acquired in a genomic background that is not currently among the dominant global *bla*_NDM_-positive lineages. Surveillance strategies that focus exclusively on dominant clones may therefore miss colonizing isolates in vulnerable wards that carry mobile resistance elements capable of horizontal spread (18–22).

Several limitations should be acknowledged. ETFK6090 was recovered from a perianal swab during colonization screening rather than from a sterile-site infection; therefore, its direct clinical significance relates primarily to colonization and transmission risk. The comparative analysis was based on publicly available complete genomes and is vulnerable to sampling, sequencing and reporting biases. The conjugation assay confirmed transferability under laboratory conditions but did not determine transfer frequency. Finally, structural modelling and docking were exploratory and should be validated by enzyme kinetics, zinc-restriction assays and, ideally, additional *bla*_NDM-15_-positive isolates. Broader prospective surveillance will be required to determine how frequently the ST580-*bla*_NDM-15_ combination occurs and whether the same IncX3 plasmid circulates in related hospital strains.

In summary, ETFK6090 represents a *bla*_NDM-15_-positive ST580 *K. pneumoniae* isolate in which carbapenem resistance was linked to a transferable IncX3 plasmid embedded within a conserved *bla*_NDM_-associated mobile context. These findings support broader genomic surveillance of carbapenem-resistant *K. pneumoniae* beyond dominant epidemic clones, particularly in high-risk paediatric and intensive-care settings.

## Ethics statement

The study protocol was approved by the Clinical Research Ethics Committee of the First Affiliated Hospital, Zhejiang University School of Medicine (FAHZU Ethics_2023-0994-Quick). A waiver of informed consent was granted by the ethics committee for the use of de-identified surveillance isolates and associated clinical information. All research procedures were performed in accordance with the ethical standards of the institutional research committee and the 1964 Declaration of Helsinki and its later amendments.

## Acknowledgement

Not applicable.

## Access to data

The whole-genome sequence of strain ETFK6090 has been deposited in the NCBI GenBank database under the accession number PRJNA1453145.

## Contribution

**Zhenghao Lou:** Writing-Original Draft, Formal Analysis. **Chengcheng Ye:** Formal Analysis, Visualization. **Xiaolu Yang:** Validation. **Qiyu Liu:** Investigation, Data Curation. **Chennuo Wang:** Project Administration, Software. **Hao Xu:** Resources. **Beiwen Zheng:** Conceptualization, Supervision, Funding Acquisition. **Xiawei Jiang:** Methodology, Writing-Review & Editing.

